# Modulation of the *Neisseria gonorrhoeae* drug efflux conduit MtrE

**DOI:** 10.1101/208504

**Authors:** Giulia Tamburrino, Salomé Llabrés, Owen N. Vickery, Samantha J. Pitt, Ulrich Zachariae

## Abstract

Widespread antibiotic resistance, especially of Gram-negative bacteria, has become a severe concern for human health. Tripartite efflux pumps are one of the major contributors to resistance in Gram-negative pathogens, by efficiently expelling a broad spectrum of antibiotics from the organism. In *Neisseria gonorrhoeae*, one of the first bacteria for which pan-resistance has been reported, the most expressed efflux complex is MtrCDE. Here we present the electrophysiological characterisation of the outer membrane component MtrE and the membrane fusion protein MtrC, obtained by a combination of planar lipid bilayer recordings and *in silico* techniques. Our *in vitro* results show that MtrE can be regulated by periplasmic binding events and that the interaction between MtrE and MtrC is sufficient to stabilize this complex in an open state. In contrast to other efflux conduits, the open complex only displays a slight preference for cations. The maximum conductance we obtain in the *in vitro* recordings is comparable to that seen in our computational electrophysiology simulations conducted on the MtrE crystal structure, indicating that this state may reflect a physiologically relevant open conformation of MtrE. Our results suggest that the MtrC/E binding interface is an important modulator of MtrE function, which could potentially be targeted by new efflux inhibitors.

## Introduction

The introduction of antibiotics into clinical use against bacterial infections marked one of the most important milestones in medicine. However, the high evolutionary pressure caused by the widespread use of antimicrobial drugs has led to the rise of antibiotic-resistant bacterial strains^1^. In recent decades, antimicrobial resistance (AMR) has evolved into a major health problem, as many bacterial species have become insusceptible to a growing range of antibiotics, and we may soon face the prospect of a post-antibiotic era. Some bacteria, especially Gram-negative organisms including forms of *Neisseria gonorrhoeae*, have become pan-resistant, i. e. they can no longer be treated with any available antibiotic^2^. The exceptional urgency of addressing the emergence of bacterial multi-drug and pan-resistance has therefore been widely recognised by national and international health authorities^3^ and *N. gonorrhoeae* has been named amongst the 12 bacteria which have been prioritised by the WHO for accelerated research efforts to develop new antibiotics^4^.

Gram-negative bacteria possess a double-membrane cell envelope, which acts as a highly efficient barrier for the inward permeation of drugs. Antibiotic agents enter these organisms predominantly via porin channels in the outer membrane, and are often expelled directly from the periplasm, located between the two bilayers, by active drug efflux pumps. Many highly resistant forms of Gram-negative bacteria display acombination of single-point mutations in porin channels and upregulation of efflux pump expression^5,6^. Amongst these, the major drivers of super-resistant phenotypes in Gram-negative bacteria are tripartite efflux pumps, protein complexes which span both the inner and outer membrane and form a continuous aqueous pathway from the inner membrane to the external medium^7^. The most clinically relevant family of Gram-negative efflux pumps is the resistance-nodulation-cell division (RND) superfamily, consisting of three major elements: an inner membrane pump protein (IMP), an outer membrane channel protein (OMP), and a membrane fusion protein (MFP) connecting the aforementioned components^8^. The importance of active efflux for the development of bacterial resistance has been impressively demonstrated in experiments, showing that multidrug-resistance can be reversed by knocking out the expression or inhibiting the function of efflux pumps^9,10^. It is therefore important to illuminate the mechanisms underpinning rapid drug expulsion. For example, targeting this major driver of multidrug resistance by new inhibitors may reduce the efficiency of efflux pumps and thereby restore the susceptibility of resistant bacteria to existing antibiotics. In addition, the modulation of efflux pumps may narrow the compound spectrum of expelled drugs and enhance the uptake of therapeutics into Gram-negative bacteria.

**Figure 1.**
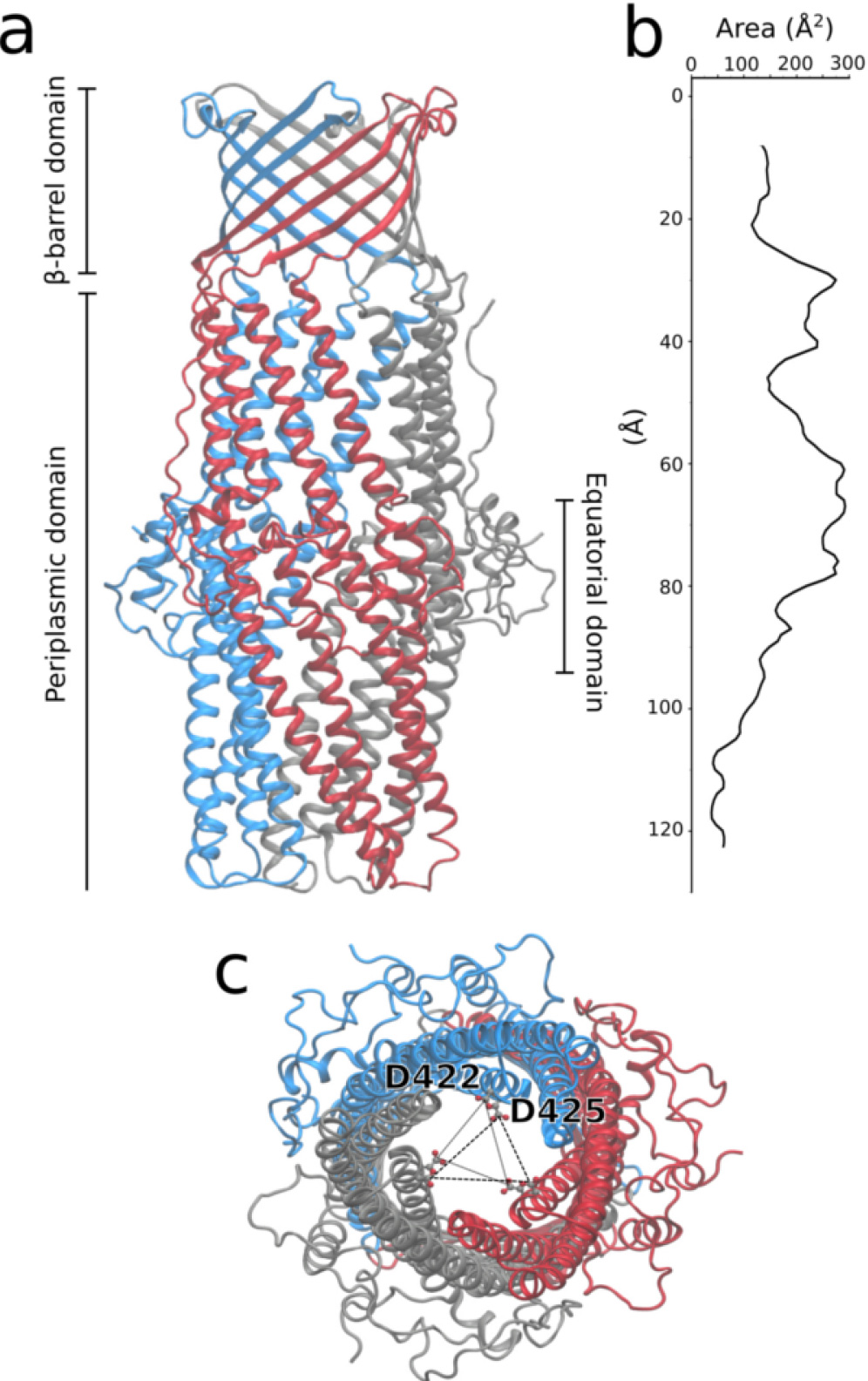
(A) Overall structure of the trimeric MtrE efflux protein channel (PDB code: 4MT0). Each subunit consists of a *²*-barrel domain,embedded in the outer membrane, an *±*-barrel domain, spanning ~100 Å inside the periplasm and an equatorial domain, mostly unstructured, located in the middle of the *α*-barrel. (B)The reported crystal structure of MtrE corresponds to a conduit with a comparably large cross-sectional area, in which the major constriction site is located at the tip of the periplasmic domain (110-120 Å from the extracellular exit along the pore axis). (C) The narrowest site in the channel is formed by two concentric aspartate rings comprising three Asp422 and Asp425 in the trimer (conserved across the other characterized OMPs), which are thought to act as a selectivity gate. Viewed from the periplasmic entrance, the ring formed by Asp422 is more proximal, the one from Asp425 more distal.

In the case of *N. gonorrhoeae*, targeting efflux is of particular urgency as to some of its strains have developed pan-resistance, making efficient medical treatment by antibiotics impossible^11^. The most highly expressed tripartite efflux pump in *N. gonorrhoeae* is MtrCDE^12^. Its outer membrane channel component, MtrE, has recently been structurally characterised (PDB code: 4MT0^13^). As in other RND homologues, MtrE is a homotrimeric protein, consisting of three domains, a *β*-barrel domain embedded in the outer membrane, an *α*-barrel domain, projecting over 100 Å into the periplasm, and an equatorial, mostly unstructured domain, located in the middle of the *α*-barrel (Fig 1a). Its putative gating region is located at the periplasmic tip and is formed by two concentric rings of conserved aspartate residues (D422 and D425; Fig 1b,c). So far, MtrE is the only structurally determined wild type RND-OMP showing an open conformation in this region, expected to maximise efflux through the duct.

Here, we present planar lipid bilayer recordings and all-atom molecular dynamics simulations characterising the conductance of the MtrE RND exit tunnel. Our results show that binding of the membrane fusion protein MtrC to the external face of MtrE stabilises the open state of the channel. Our findings thus highlight the contact region between the OMP and the MFP of MtrCDE as a switch between open and closed states of the outer drug efflux conduit. This interface may therefore represent an attractive site for potential molecular intervention.

## Results

### Planar Lipid Bilayer Experiments

First we performed electrophysiological measurements, in which MtrE was embedded into symmetric POPE planar lipid bilayers. Planar lipid bilayer electrophysiology has frequently been used to investigate the conductance and gating of OMPs in response to membrane voltages^14,15,16,17^. The method makes use of simplified membrane models, as opposed to the highly complex architecture of bacterial outer membranes, which consists of asymmetric lipid bilayers, with inner leaflets containing phospholipids and outer leaflets composed mainly of lipopolysaccharides^18^. It has however been shown that electrophysiology in these simplified membrane models is a valid approach to characterise the translocation of ions across outer membrane channels^19^. Furthermore, the gating region of efflux outward gates is located at a large distance from the membrane in the periplasmic space, and therefore it is expected that the simplified membrane does not substantially affect results on the gating of these outer conduits^16^.

It has further been demonstrated that the direction of insertion of OMPs into membranes is entirely determined by the protein structures^20,21^and that OMPs spontaneously insert unidirectionally in planar lipid bilayers^16^. In addition, MtrE possesses a large polar periplasmic domain, which is unlikely to traverse the membrane (Fig 1a). Since we have exclusively added the OMP to the membrane face corresponding to the periplasmic side (cis-chamber; n=10), it is highly probable that MtrE is unidirectionally inserted into the bilayer in our experiments, despite the symmetry of the POPE leaflets.

When voltage-clamped at +40 mV, the fully open state of MtrE was characterised by a unitary current amplitude of 11.5 pA (Fig 2A). MtrE was also found to open to multiple sub-conductance open states that were approximately 18% (2 pA) and 60% (7 pA) of the fully open state (Fig 2A). Addition of the binding partner MtrC to the cis-chamber significantly increased the open probability of MtrE from 0.35±0.11 to 0.86±0.12 (n=3) and caused MtrE to gate predominantly to the fully open state (Fig 2A). Transitions to the sub-conductance open states of MtrE following the addition of MtrC were no longer resolved.

When only a single MtrE channel was gating in the bilayer, lifetime analysis revealed that in the absence of the MtrC membrane fusion protein MtrE displayed fast flickery gating (Fig 2B). The addition of MtrC to the periplasmic face of MtrE altered channel gating causing the channel to dwell for longer sojourns in the fully open state, demonstrated by an increase in the apparent open time from 1 ms to 10.5 ms (Fig 2B). This suggests that MtrC alters the conformation of the MtrE protein and stabilizes the channel in the fully open state. In 3 out of 13 experiments, MtrE gated predominantly to the fully open state even in the absence of MtrC adapter protein, suggesting that once in this state, channel openings are stabilised. Construction of a current-voltage relationship for the fully open state of MtrE displayed an open channel conductance of 304±7 pS (Fig 3A).

Under non-symmetrical conditions (trans 210 mM KCl:cis 460 mM KCl), the reversal potential shifted to −5.5±0.8 mV (Fig 3B), revealing that MtrE is approximately 1.7 fold more permeable for K^+^ than for Cl^−^. This characterises MtrE as a slightly cation selective channel. The best-studied OMP homologues, TolC from *E. coli* and OprM from *P. aeruginosa*, also display a preference for cations, however to a much larger degree. For instance, TolC has been found to be 16.5 fold more permeable for K^+^ than for Cl^−^^22^, and the selectivity of OprM has been estimated to be similar to that of TolC^23^.

**Figure 2.**
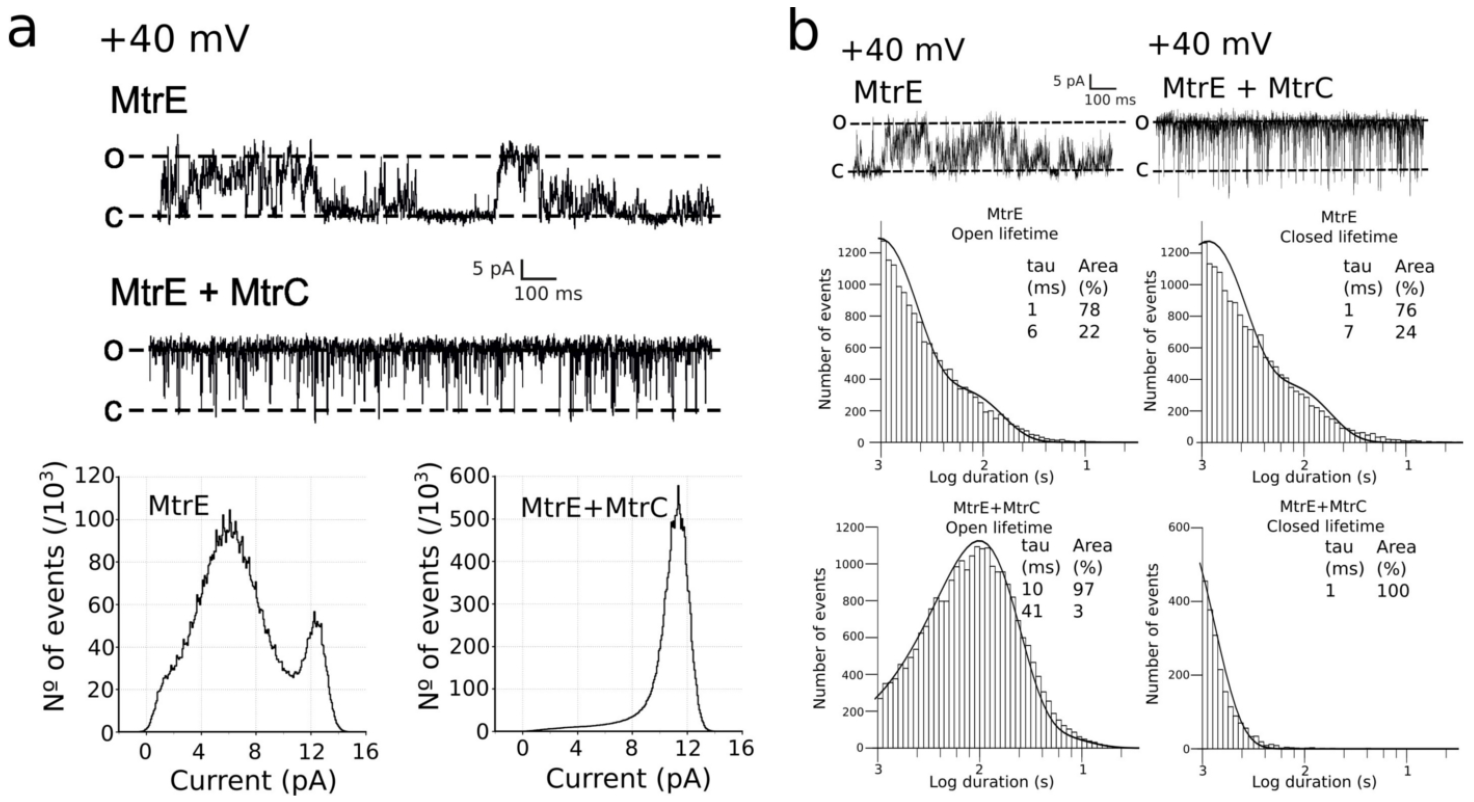
(A) Single channel experiments reveal that MtrE can dwell between 3 different semi-open states. The addition of the adapter protein MtrC stabilizes the most open conformation. (B) Lifetime analysis uncovers that the effect of MtrC is to significantly reduce the closed lifetime of MtrE, so that the channel resides for shorter intervals in the closed state.

**Figure 3.**
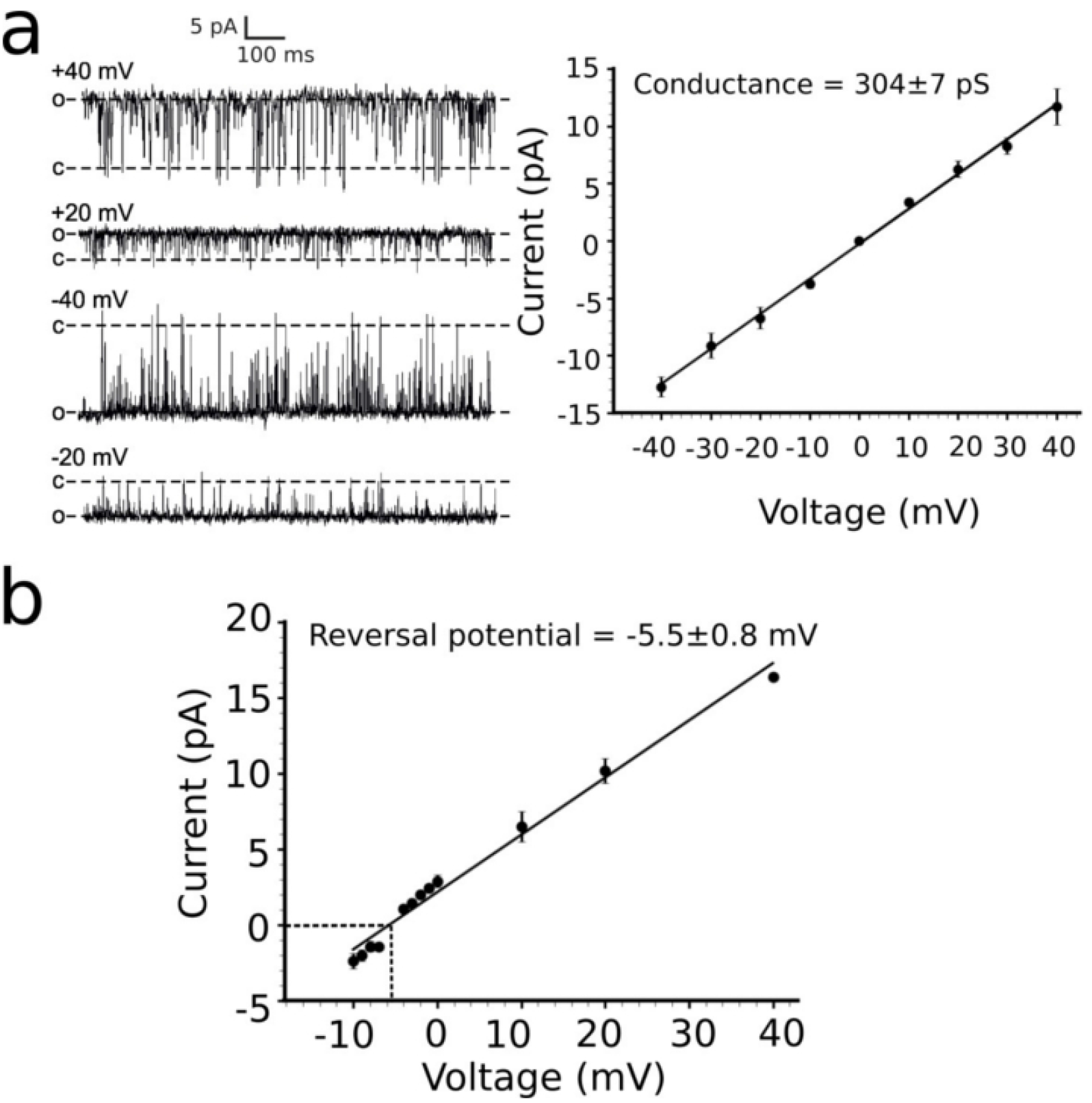
(A) The I/V plot of the MtrC/E protein complex is linear in the experimental voltage range (between −40 and +40 mV) and the conductance for the fully open state is 304±7 pS. (B) The reversal potential of MtrE in non-symmetrical conditions (trans 210 mM KCl:cis 460 mM KCl), indicated by a dotted line, is at −5.5±0.8 mV revealing slight cation selectivity.

### MtrE Dynamics and Computational Electrophysiology

Previous experiments on the cellular level have shown that substrates cannot traverse MtrE from the extracellular space to the periplasm when complex formation to the tripartite pump is inhibited^24^. This finding suggests that MtrE requires the interaction with other pump components to induce the fully open conformation, which is expected to maximise the efficiency of efflux. In the crystal structure, the Asp gating rings (Fig 1c) and the overall conformation of MtrE reflect an open state, wide enough to accommodate even larger substrate molecules (Fig 1b), although its complex binding partners are absent. We were therefore interested if the crystallographically observed conformation corresponds to the fully open state of MtrE.

**Figure 4.**
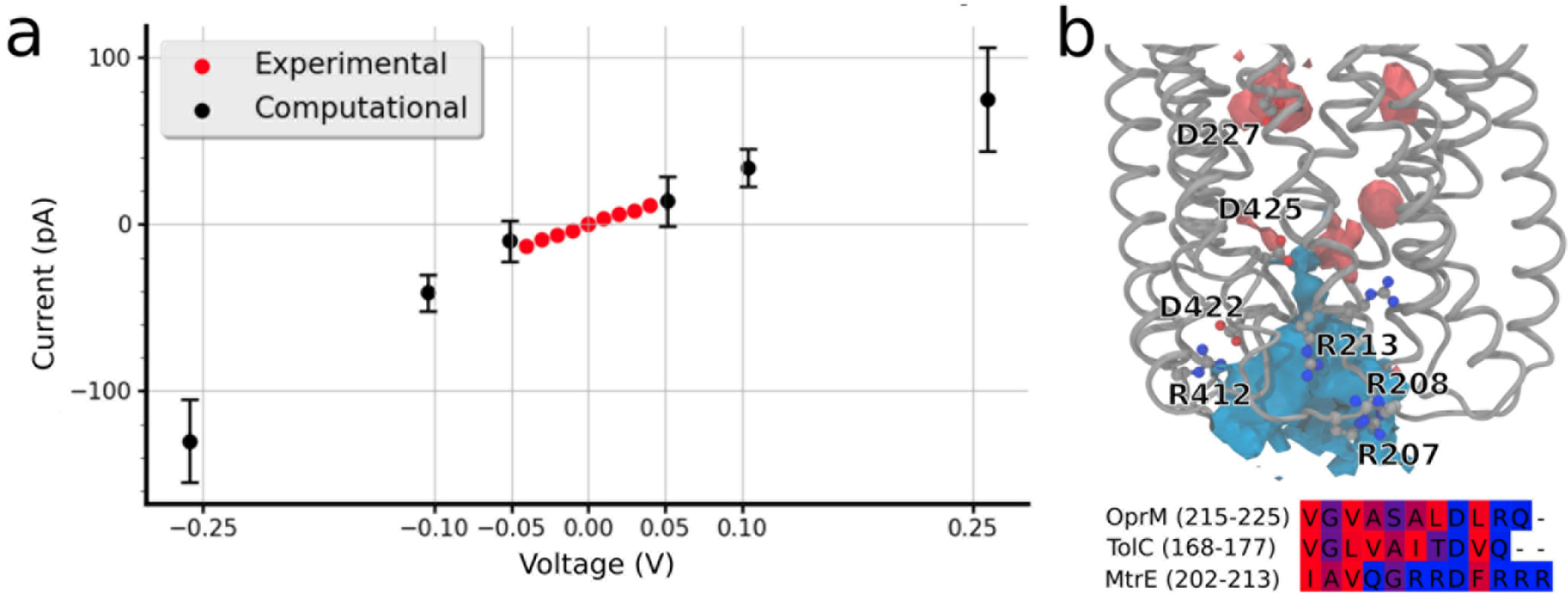
(a) Red dots display the experimental current-voltage relationship in symmetrical 210 mM KCl solution, black dots show the current/voltage relationship obtained from the MD simulations near the experimental voltage range. Voltages of ±0.05, ±0.10 and ±0.25 V were applied along the z-axis using external electric fields as implemented in GROMACS^25^, together with the charmm36 force field and positional restraints of 200 kJ/mol nm^2^ on MtrE heavy atoms as detailed in the text. We observe excellent agreement between the experimental and the computational currents. The computational x-axis and the experimental x and y-axis error bars are smaller than the data points. (b, bottom) Sequence alignment of the three electrophysiologically characterised efflux OMPs. MtrE shows an abundance of arginine residues in the periplasmic loop region in addition to a series of aspartate residues lining the interior of the pore, which is not present in homologous proteins. (b, top) The density map of K^+^ (contoured at 2% SD, in red) and Cl^−^ (contoured at 5% SD, in blue) highlights the influence of this highly basic region on ion flow through the channel.

The majority of our simulations of MtrE immersed in POPC and POPE membranes show rapid closing of the periplasmic gating region. Despite displaying complex dynamics, especially in the loop region and the outer helices, the inner helices rapidly adopt a continuously closed state on the time-scale of most of our simulations. This finding indicated that the protonation state of the Asp gating rings may act as a potential modulator of the closing transition^26^. We therefore tested if a different protonation state of the Asp side chains affected channel closing. As shown in supplementary Fig S1, protonation of the Asp gating rings slightly lowers the propensity for closing, however a consistently stable open state of the gating region is still not observed. This observation suggests that additional factors are likely to play a role in stabilising the open state of the channel. Our experimental data show that binding of MtrC leads to the stabilisation of MtrE in an open conformation for substantial lengths of time. Taking these results together, we suggest that binding partners are required to retain MtrE in a fully open conformation, while protonation changes within the gating region, possibly induced by binding, may further stabilise the open state. In addition, our results indicate that crystal contacts may be able to at least partially compensate for the lack of interactions with other pump components to retain MtrE in the open state. In our single channel experiments, we observed MtrE to adopt a mostly closed state in the absence of MtrC, while the pH of the solution was not altered, emphasising the major role of the MtrE/MtrC contacts we find for gating the channel.

To date, there is no experimental information available on the three-dimensional structure of MtrC, nor its mode and position of binding to MtrE. To mimic the stabilising effect of the membrane fusion partner, we therefore restrained the overall backbone conformation of the protein and investigated by computational electrophysiology simulations (CompEL) if the pore geometry captured in the crystal can explain the experimentally observed channel currents. Due to the limited time-scale of the simulations, we used slightly raised transmembrane voltages compared to our experimental conditions, in order to drive ion permeation and ensure sufficient sampling^27^. Fig 4 shows that the current we obtain for single open MtrE conduits is generally in good agreement with our experimental data. The estimated maximum conductance of the open pore is 324±34 pS, which is similar to the maximum experimental conductance of 304±7 pS. In the simulations, we observe ion selectivity ratios between 1:1.2±0.5 (K^+^:Cl^−^, at 100mV) and 1:1.3±0.5 (at -100mV). These voltages were chosen as they are close to the experimental range of voltages, while at the same time allowing us to record sufficient sampling of ion permeation for robust selectivity estimates. The selectivity values are in generally good agreement with the experimental ion selectivity, although Cl^−^ ions show a slightly higher permeability in our simulations compared to the experiments.

We modelled an additional MtrE conformation, based on the most dilated state observed for TolC so far, where it is in complex with the complete RND machinery MacAB-TolC^28^ (Supplementary Fig S2). For this model, we measure an *in silico* conductance of 597±70 pS, which is considerably higher than our experimental conductance. Our data therefore suggests that the crystal structure of MtrE is a good representation of the overall conformation of the fully open state of this conduit. Our results further show that protein-protein contacts between MtrE and MtrC stabilise the fully open pore in the electrophysiological experiments, whereas in isolation, such as in our computer simulations, the open conformation of MtrE is likely to undergo rapid transitions to more closed states unless restrained. Supporting this notion further, we constructed a molecular model of the bound state of MtrE and MtrC, based on the homologous complex of AcrAB-TolC resolved by electron cryo-microscopy, which exhibits a similar degree of opening of the OMP gating region as the MtrE crystal structure^8^. Simulations of this complexed model indicate an increased tendency to remain in an open state without undergoing further protonation changes (Fig S3).

Our analysis of ion trajectories in the simulations of MtrE under voltage (Fig 5) shows that, although both anions and cations generally occupy most parts of the MtrE channel lumen to similar extent, there are regions in which important differences are observed. Especially the periplasmic gating region, but also sections of the transmembrane *β*-barrel show a substantially reduced K^+^ density, particularly evident at negative membrane potentials (Fig 5a,b, left). A superposition of the positions of ions along the trajectories shows the constrictions observed for the cation pathway at the periplasmic entrance, highlighting the importance of this gating region, which includes a high density of arginine residues (see Fig 4), for the control of ion conduction. Furthermore, we observe that the channel lumen and gating region are well-hydrated in the open state of MtrE (Fig 5a,b, right).

Dewetting transitions play a major role in the gating process of many membrane channels^29,30^. To ascertain if dewetting plays a part in the closing of MtrE, we further analysed the simulations of MtrE, in which the channel rapidly adopted a closed state (Fig S1). We find that even when the gating region shows a minimal level of openness, occupation with water and a continuous water chain between the extracellular and periplasmic side are still observed (Fig S1B). Our findings suggest that dewetting of MtrE is unlikely to underpin the gating of this channel for ion conduction.

**Figure 5.**
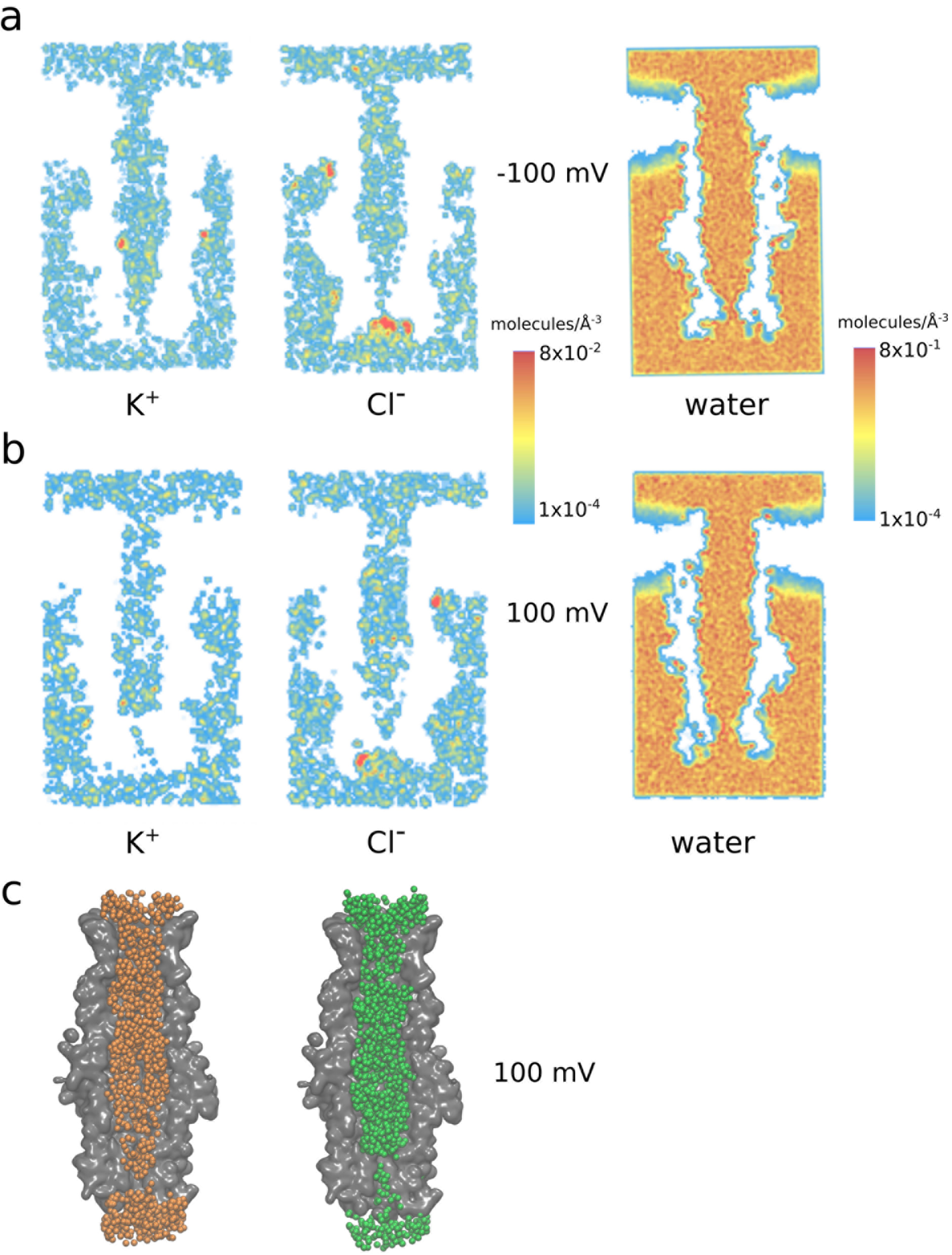
(a-b) Pathways of K^+^ and Cl^−^ ions in the MtrE channel (depicted as ion density) and average water density in the channel during simulations at (a) +100mV and (b) −100mV. The panels show the extracellular side of MtrE at the top and the periplasmic gate at the bottom of each figure. Under both conditions, a thinning out of the ion density at the periplasmic gate can be observed, especially for K^+^ ions. (c) Superposition of all K^+^ (orange) and Cl^−^ (green) ions during 250ns (1 frame every 2.5ns, 100 total frames) at +100mV clearly shows this constriction of the ion pathways within the gating region, representing the major bottleneck for ion permeation.

## Discussion

Both our experimental data, obtained from single-channel electrophysiology, and our computational results suggest that the channel MtrE is mainly present in a closed state when it is unbound from the rest of the efflux machinery. These findings are in agreement with electrophysiological recordings on homologous OMPs^16,31^and biochemical data on MtrE^32,33^. Our results show that the interaction of the adapter protein MtrC with MtrE is sufficient to stabilise the complex in an open state, which conducts large ion currents in accordance with the cross-section of the channel observed in the MtrE crystal structure^13^. This is, to our knowledge, the first time that the fully open state of an OMP from an efflux pump could be characterised without inserting mutations in the gating region, as previously reported, e.g., for TolC from *E. coli*^31^.

We attribute the subconductance states we observe in the electrophysiological recordings to the range of closed conformations we find in the MD simulations, which show a remaining open channel cross-sectional area of about 30-40Å^2^ and continuous hydration of the pore (Fig S1). As opposed to TolC, which shows a strong preference for cations^22^, the cation selectivity we observe for MtrE is about 10-fold smaller. In the simulations, a region of diminished cation density in the channel coincides with an abundance of arginine residues in the loop region at the periplasmic tip of MtrE, which precedes the channel-internal conserved aspartate gating rings from the periplasmic side, suggesting a role for these arginine groups in determining the ion selectivity of MtrE (Figs 4 and 5). Ultimately however, additional experiments will be required to unequivocally identify the determinants of the lowered level of MtrE ion selectivity. Notably, the major outer membrane porins of *Neisseria*, PorA and PorB, and of related organisms (e.g. Omp32 from *Delftia acidovorans*), show strong anion selectivity^34,35,36^in contrast to the cation-selective porins of other Gram-negative organisms (such as OmpF from *E. coli*)^37^. The reduced cation-selectivity of *N. gonorrhoeae* MtrE could thus be linked to an increased inward permeability of the outer membrane of *Neisseria* species for anions compared to other Gram-negative bacteria.

All of these previous findings, together with our results, suggest that the inward and outward permeability of neisserial outer membranes differs substantially from that of model organisms, frequently used to investigate the determinants of Gram-negative cell wall permeation, such as *E. coli*. In particular, the uptake and efflux of antibiotics may be underpinned by different principles. According to our data, the effect of different model lipids on MtrE function is rather small, but the complex membrane composition of neisserial outer membranes may add a further layer of modulation and control of neisserial OMPs, which may differ from observations made in other Gram-negative bacteria.

The differences to permeation across *E. coli* TolC, for example, may have important consequences for the design of antibacterials against these difficult-to-treat Gram-negative pathogens. In particular, *N. gonorrhoeae* strains are amongst the most antibiotic-resistant bacteria, displaying pan-resistance against all presently available antibiotic agents^11,4^. This confers a high degree of urgency to the development of novel therapies against *N. gonorrhoeae* infections.

Active drug efflux of a broad spectrum of antimicrobials is one of the major factors driving the development of resistance in Gram-negative bacteria^2^. Our results suggest that the less stringent selectivity for cations in MtrE we found here may contribute to the efficiency with which a very wide range of antibiotics is expelled from *N. gonorrhoeae* by the MtrCDE efflux system. The diminished cation-selectivity may however also allow the exploration of new chemical space for the design of clinically usable efflux inhibitors. Most previous attempts at designing efflux pump inhibitors have failed in clinical studies due to toxicity problems related to the cationic pharmacophore, which is required for efficient competitive inhibition of the pumps^38^. Some inhibitors were also targeted against the internal Asp gating rings of the OMP, but clinical success was not achieved by these cationic inhibitors either^39^. The different electrostatics of the MtrE interior, and its reduced preference for cations, may enable drug researchers to design novel inhibitors with different charge properties.

Importantly however, our results indicate that it may not be necessary to block the outer gate of the efflux system through orthosterically binding inhibitors. We show that the opening of MtrE is regulated by allosteric binding events of the adapter protein MtrC on the periplasmic outer face of the pore. Association with MtrC alone is sufficient to keep MtrE in a prolonged open state. The binding of MtrC may be linked to further protonation changes at the interface and pore interior of MtrE. These findings suggest that the MtrE-MtrC binding interface may be an attractive targeting site for the development of allosterically acting efflux inhibitors. These inhibitors would no longer compete for the orthosteric drug binding sites in the efflux pumps, but still regulate the openness of, and thereby the efficiency of drug expulsion across, the outward conduit. This new modulation mechanism may potentially facilitate the design of a new type of inhibitor, avoiding previous chemotypes known for their toxicity.

## Methods

### Computational methods

The 3.29-Å resolution crystal structure of MtrE (PDB code: 4MT0)^13^ was used as a starting structure. The sulfate ion was removed and the N- and C-terminal residues were capped using acetyl and N-methyl amide groups, respectively. We note that there is relatively weak electron density for residues 203−212 in the crystal structure. All atomistic molecular dynamics simulations were performed using the GROMACS-5.1.1 software package^40^. We tested the robustness of our results by using different forcefields.

First, the protein was embedded in pre-equilibrated and solvated POPC (1-palmitoyl 2-oleoyl sn-glycero 3-phosphatidyl choline) bilayers containing 288 lipid molecules (177 after insertion of the protein) using the GROMACS g_membed utility^41,42^. The system was solvated and K^+^ and Cl^−^ ions were added to neutralise the system and to reach a concentration of 200 mM. Here, the Amber ff99SB-ILDN force field was used for the protein^43,44^, and Berger parameters adapted for use within the Amber ff99SB-ILDN force field were employed for the lipids^45,46^. The SPC/E model was used for the waters^47^ and Joung/Cheatham III parameters were employed for the ions^48^. The systems were minimised and then equilibrated with position restraints on protein heavy atoms of 1000 kJ/mol·nm^2^ for 20 ns. Water bond angles and distances were constrained by SETTLE^49^, while all other bonds were constrained using the LINCS method^50^. The temperature was kept constant at 310K, using the v-rescale method^51^ with a time constant of 0.2 ps. The pressure was kept constant throughout the simulations at 1 bar, using a Berendsen barostat^52^ with semi-isotropic coupling. The application of the virtual site model for hydrogen atoms^53^ allowed the use of a 5-fs time step during the simulations. For the CompEL simulations, the protocol described by Kutzner et al.^27^ was used; the system was duplicated along the z axis to construct a double bilayer system, and ionic imbalances from 2 to 6 Cl^−^ ions were used between the aqueous compartments to generate a range of transmembrane potentials from ~-500 to ~500 mV. Some simulations made use of protein backbone heavy atom position restraints of 200 kJ/mol·nm^2^ or 1000 kJ/molnm^2^ as indicated in the text.

Furthermore, in additional sets of simulations, we employed the CHARMM36 force field for the protein, lipids and ions^54^. For water molecules, the TIP3 model was used^55^. The temperature was kept constant at 310K, using the Nose-Hoover method^56^ with a time constant of 0.1 ps. The pressure was kept constant throughout the simulations at 1 bar, using a Parrinello-Rahman barostat^57^ with semi-isotropic coupling. Here, a 2-fs time step was used, and the proteins were inserted in a POPE (1-palmitoyl 2-oleoyl sn-glycero 3-phosphatidyl ethanolamine) membrane. A constant electric field was applied to generate membrane potentials from ~-250 to ~250 mV^25^. The systems were constructed by using the CHARMM-GUI webserver^58^. In the production simulations, protein backbone heavy atom position restraints of 200 kJ/mol·nm^2^ were applied.

Structural alignment was performed using the Jalview suite^59^, together with the Clustal W program^60^. Homology modelling was performed with the MODELLER9.16 suit^61,62^, using the dilated Cryo-EM structure of TolC^28^(Fig S2). During the modelling process, symmetry restraints were applied on MtrE C*α* atoms, to take into account the threefold symmetry of the protein.

### Protein production

The MtrE protein was produced based on a previous protocol by Lei et al.^13^, but using the different expression plasmid pET28a. It was solubilised from the membrane in n-dodecyl-*β*-D-maltoside (DDM). The MtrC protein was produced based on a previous protocol by Janganan et al.^33^, increasing the growth period to 18 hours to yield a sufficient amount of protein. Analytical gel filtration suggested that the protein runs in several multimers, mainly dimers and hexamers, which correlates well with the protocol used^33^. The purity of the samples was confirmed by SDS-PAGE, and their identity was confirmed through peptide mass fingerprinting, conducted by the FingerPrints Proteomics Facility of the University of Dundee.

### Conductance measurements and analysis

Current recordings were monitored under voltage-clamp conditions using a BC-525C amplifier (Warner Instruments, Harvard) following established methods^14,63^. Recordings were low-pass filtered at 10 kHz with a 4-pole Bessel filter, digitized at 100 kHz using a NIDAQ-MX acquisition interface (National Instruments, Texas, USA). Data were recorded to a computer hard drive using WinEDR 3.6.4 (Strathclyde University, Glasgow, UK). The recordings were subsequently filtered at 800Hz(−3 dB) using a low-pass digital filter implemented in WinEDR 3.6.4. Experiments were performed at room temperature (20-22 °C). The MtrE protein was added to the cis chamber and incorporations made at +40mV, under continuous stirring. Unless otherwise stated, single-channel events were recorded in symmetrical 210 mM KCl. For selectivity measurements, the cis-chamber contained 510 mM KCl and the trans-chamber contained 210 mM KCl. Selectivity was computed using the Goldman-Hodgkin-Katz equation. E_rev_ was taken as the voltage at which no current flow was detected and was corrected for junction potential. Lifetime analysis was performed using TAC and TAC-fit software (Bruxton, Seattle, WA).

## Acknowledgements

We thank Sharon Shepherd, Thomas Eadsforth and the Protein Production Unit of the University of Dundee for expression and purification of the MtrC and MtrE proteins. We are grateful to Gavin Robertson and Ben Reilly O’Donnell for assistance with the electrophysiology measurements. We acknowledge funding through the Wellcome Trust Interdisciplinary Research Funds (grant WT097818MF) and Tenovus Tayside (grant T16/30). ONV has been funded through a BBSRC CASE award (BB/J013072/1).

### Author contributions statement

S.L., S.J.P. and U.Z. conceived and designed the research, G.T. conducted the research, all authors analysed the data, G.T., S.J.P., and U.Z. wrote the manuscript, and all authors edited and reviewed the manuscript.

---

**Figure S1.**
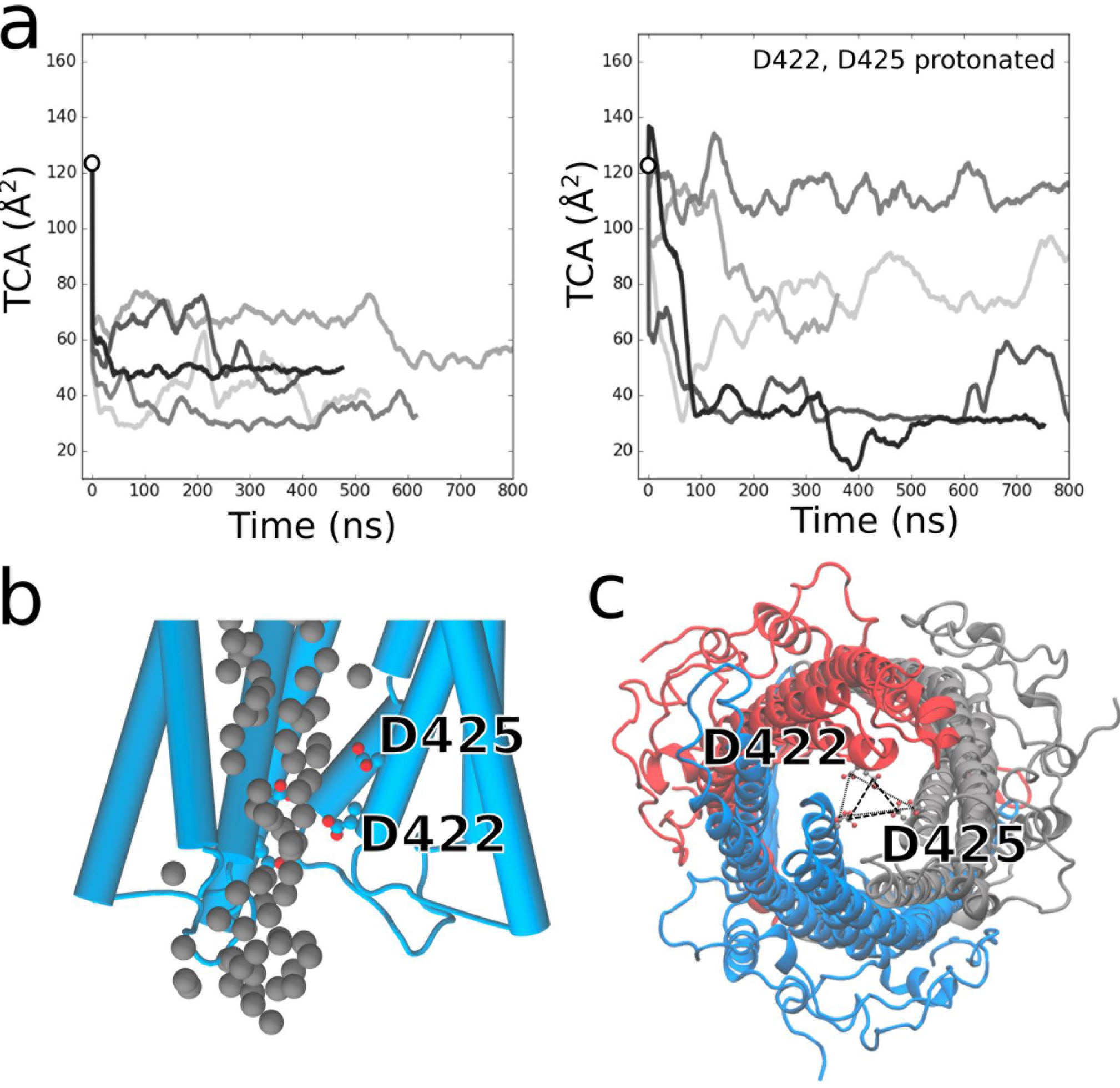
(a) We used the TCA (triangular cross-sectional area) spanned by the C*α* atoms of the Asp422 residues in each MtrE monomer to obtain a simple estimate of the cross-sectional area of the periplasmic gate. The black circle indicates the TCA of the crystal structure. In simulations where no acidic residue is in a protonated state, we observe fast closing, with a closed state that remains stable on the time-scale of the simulations (left). In simulations with uncharged protonation aspartate gating rings, we observe an effect on the closing propensity of the channel, due mainly to the disruption of essential intra extra-chain interactions, but a strong tendency to adopt a more closed state is retained in most simulations (right). (b) Representative snapshot of a closed conformation in the gating region with water oxygen atoms in gray. A continuous water wire is consistently observed inside the pore despite its closed state, and dewetting is not observed. (c) Representative closed conformation of MtrE and its TCA.

**Figure S2.**
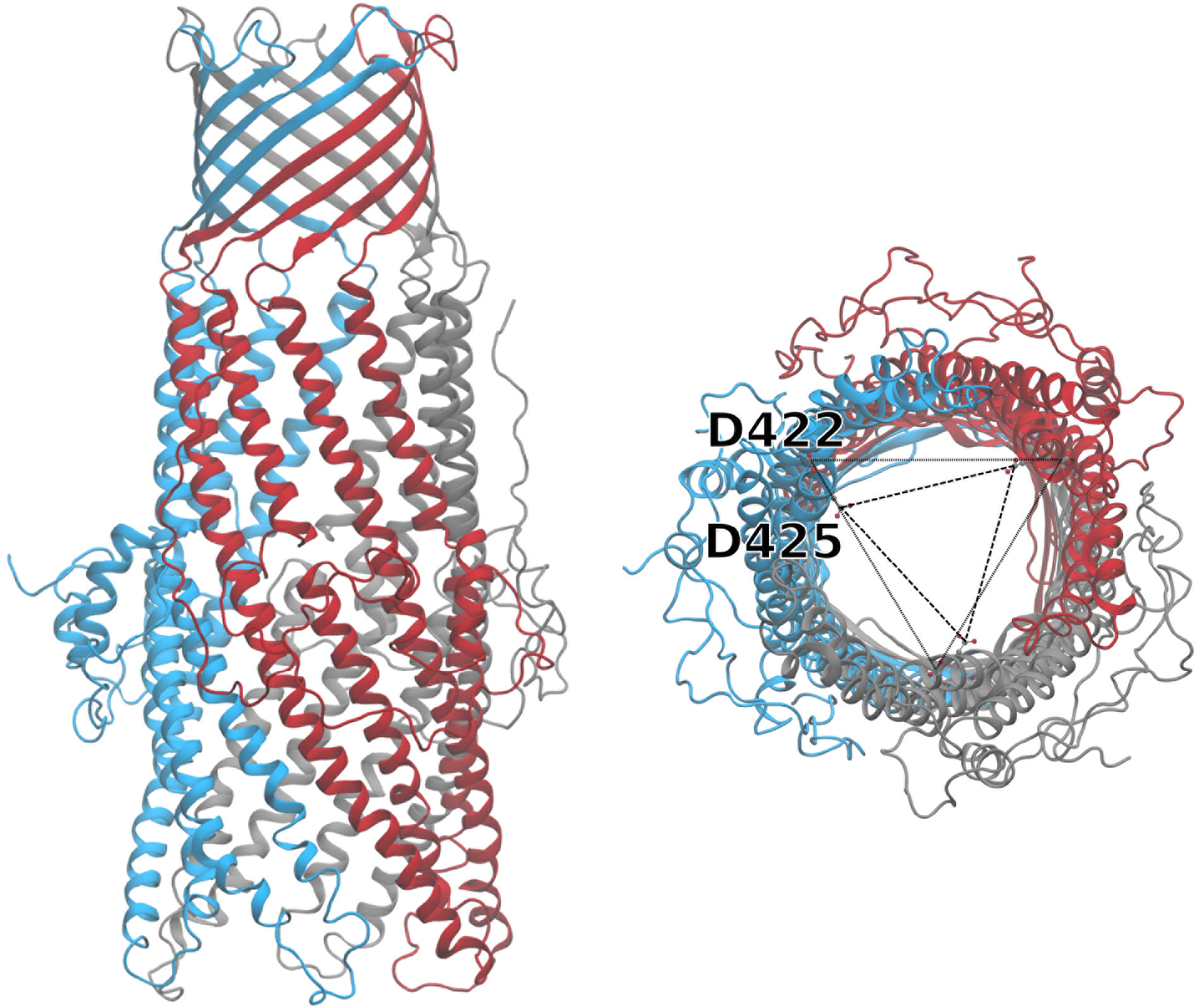
Model of a super-open state of MtrE based on the dilated conformation observed for TolC in the TolC-MacA complex from *E. coli*^28^. In our simulations, we observe a greatly increased conductance of MtrE in this dilated conformation, which is in disagreement with our experimental recordings on the channel.

**Figure S3.**
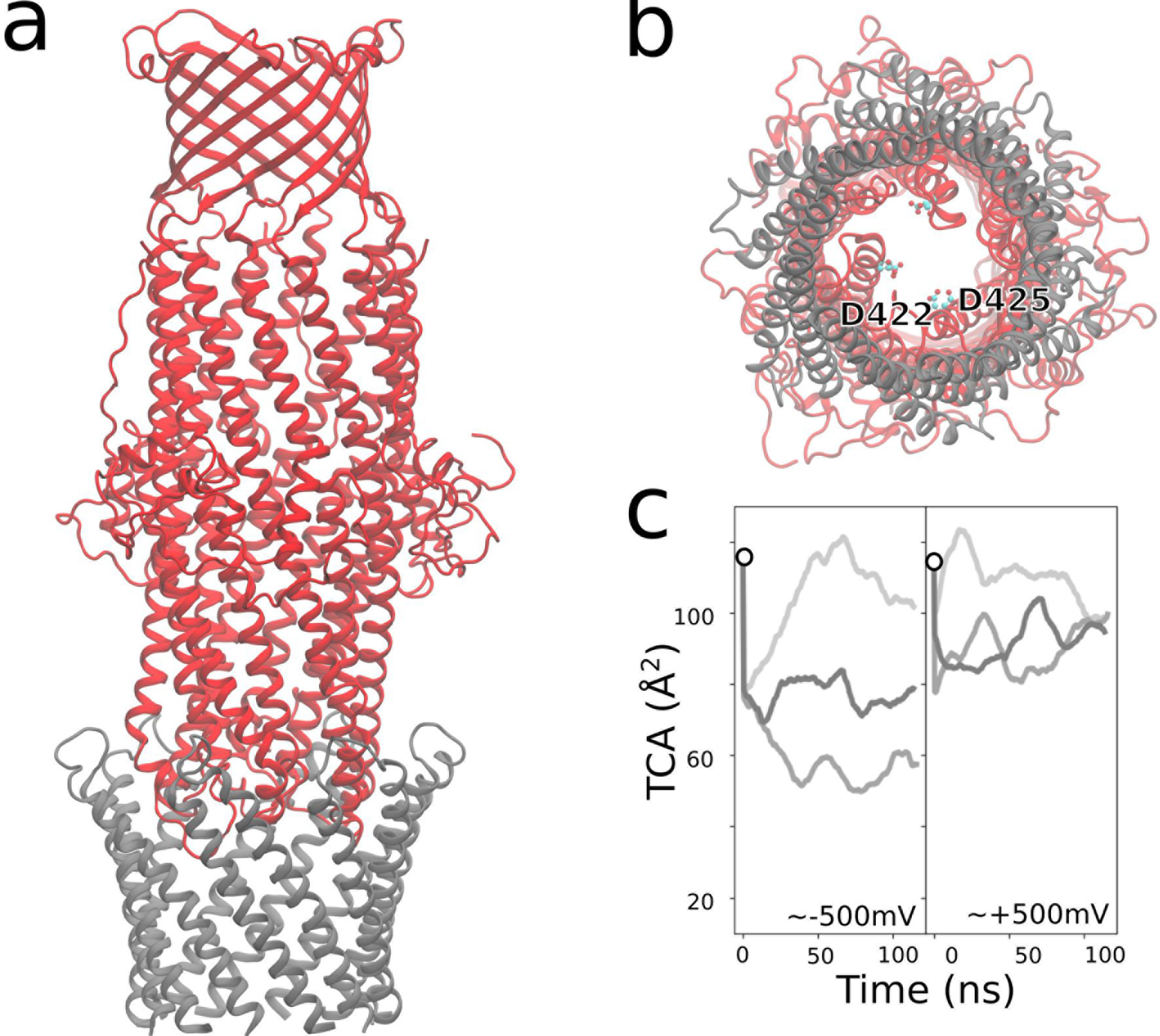
(a-b) Model of a semi-open state of MtrE in complex with the hairpin portion of MtrC, based on a conformation observed in the TolC-AcrAB complex from *E. coli*^8^ (a, side view, b, close-up view of the interface region from MtrC). The TCA of this state is similar to the TCA of MtrE in its open-state crystal structure^13^, indicated by black circles. (c) TCA of the modeled complex in simulations under voltage.

**Figure S4.**
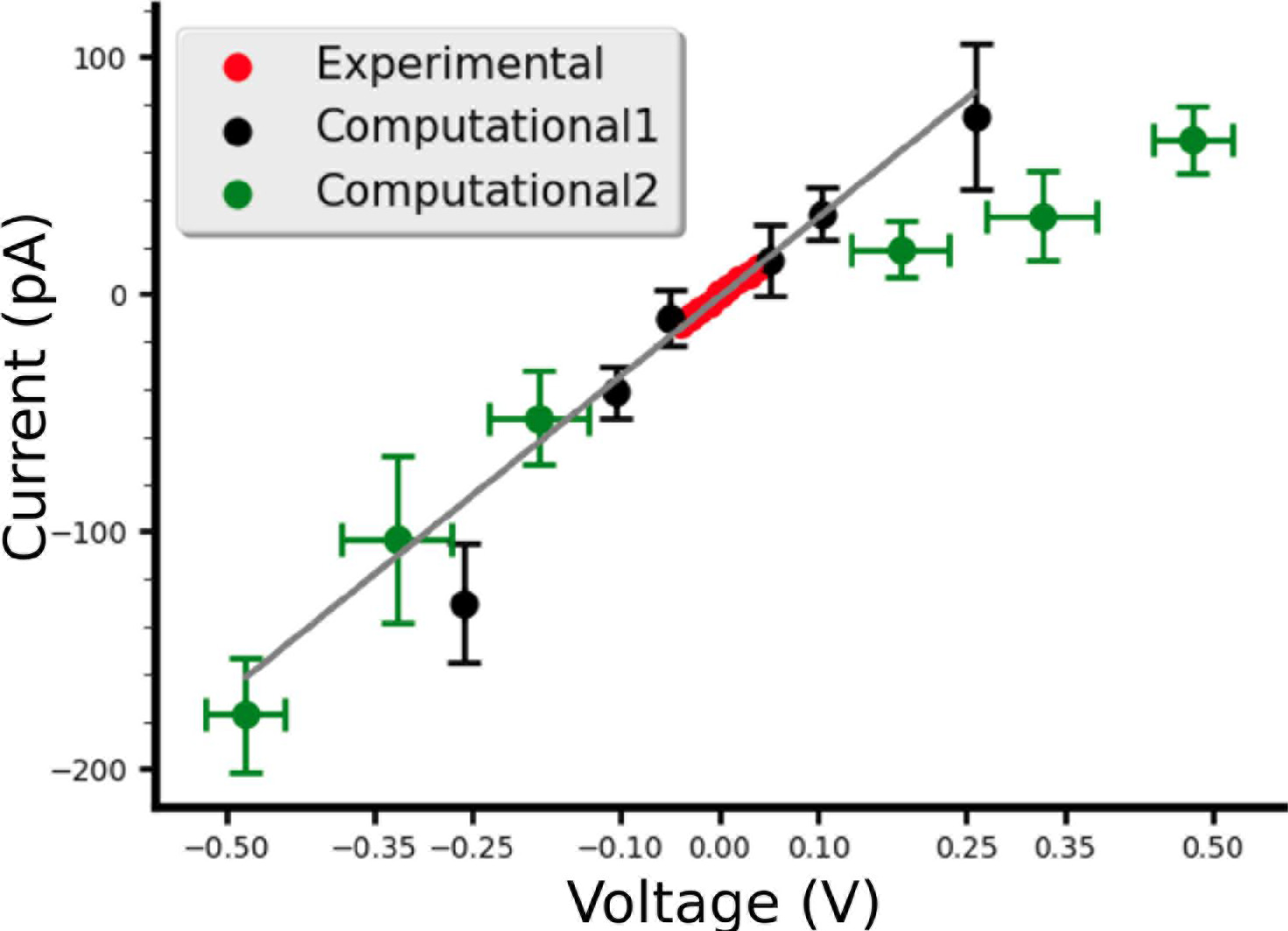
Current-voltage relationship of MtrE from experiment and simulations including voltages far from the experimentally applied ones. Red dots display the experimental current-voltage relationship data in symmetrical 210 mM KCl solution. Black dots show the current-voltage relationship obtained from MD simulations using an external electric field, green dots display data points obtained from CompEL. In the CompEL simulations, voltages of ±0.18, ±0.32 and ±0.48 V were applied by maintaining an ion imbalance of 2, 4, or 6 Cl^−^ ions, respectively, between two separate compartments. Extrapolating the linear current-voltage relationship obtained in the experimental measurements towards markedly higher voltages (gray line) suggests that at highly positive transmembrane voltages, the data points obtained from an applied electric field offer a better prediction of the MtrE conductance, while by contrast, in the highly negative voltage range, the values from CompEl more closely follow the slope of the extrapolation. Since these voltages are outside of the experimental range, the actual magnitude of the MtrE currents at these potentials remains unclear. Initial analysis shows that the difference arises due to the two distinct ways in which the transmembrane electric field is modelled, and is not related to the force-fields or the membrane lipids used. The emergence of differences between the two approaches is interesting and, since it is out of the scope of the present work focusing on MtrE modulation, will be investigated in the context of a future study.

**Figure S5.**
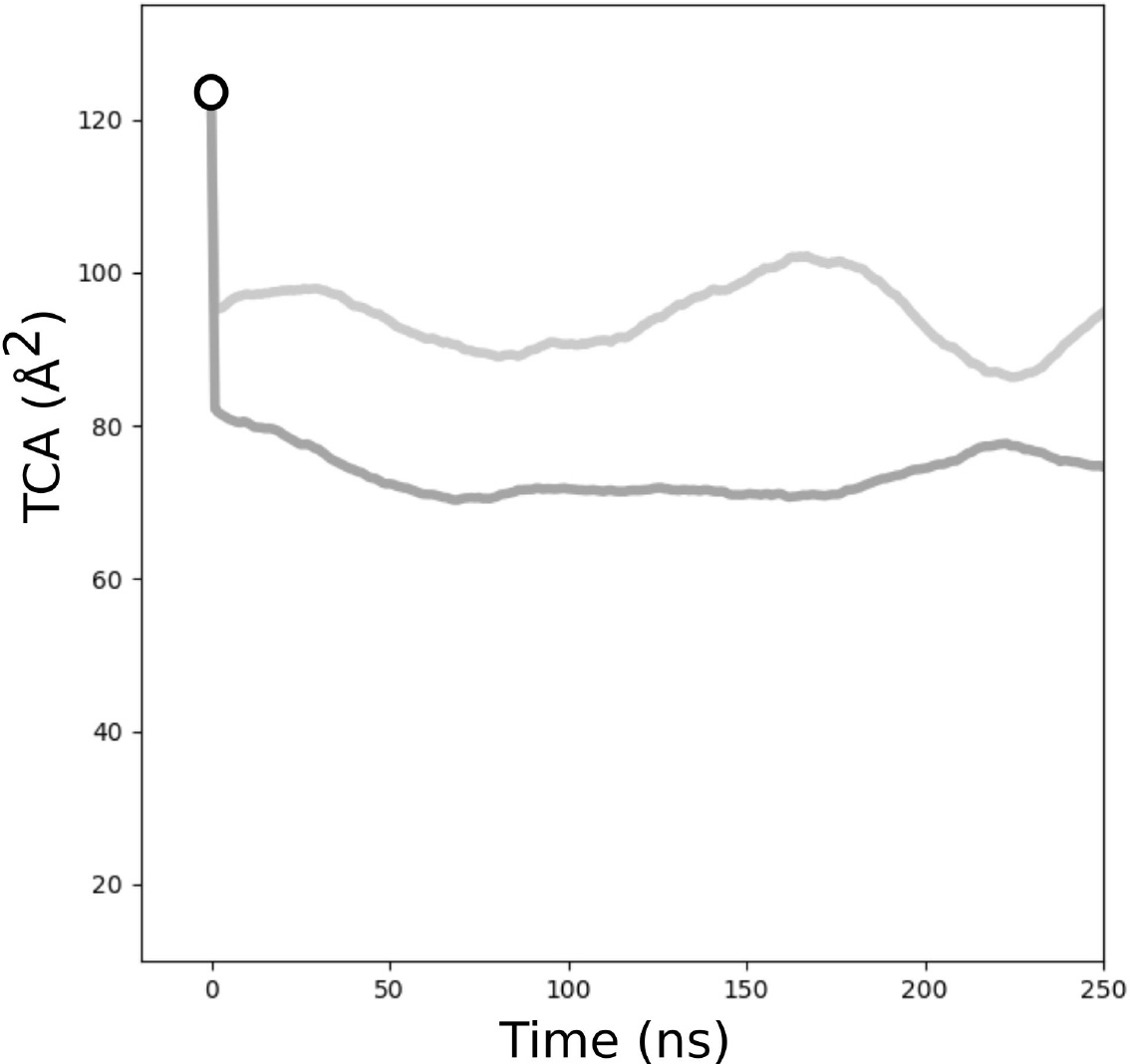
TCA spanned by the C*α* atoms of the Asp422 residues in each MtrE monomer from unrestrained simulations of MtrE in a pure POPE bilayer. The area exhibits a rapid transition of the channel to a more closed conformation compared to the crystal structure, which is 130Å^2^. This is in line with our observations made in POPC bilayers.

**Figure S6.**
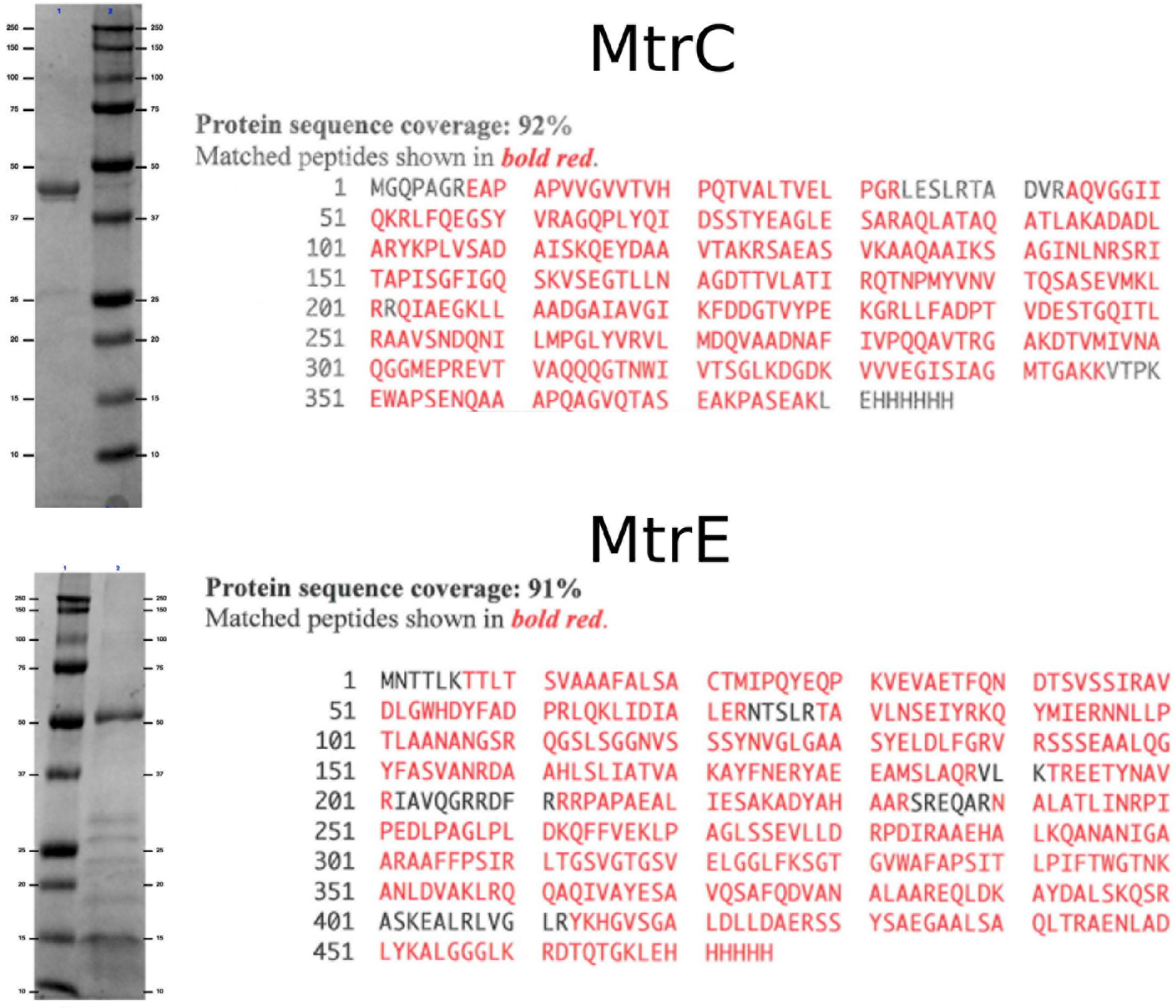
SDS PAGE and fingerprint mass-spectrometry (MS) sequence analysis for the MtrC (top) and MtrE (bottom) proteins. The MtrC sample (lane 1, top) has been purified to 94.2%, while the membrane protein MtrE (lane 2, bottom) is 50.8% pure. Both samples were confirmed as MtrC and MtrE, respectively, by MS identification.

## References

1. Neu, H. C. The Crisis in Antibiotic Resistance. Science 257, 1064–1073 (1992). DOI 10.1126/science.257.5073.1064.

2. Unemo,M. & Nicholas, R. A. Emergence of multidrug-resistant, extensively drug-resistant and untreatable gonorrhea. Future Microbiology 7, 1401–1422 (2012). DOI 10.2217/fmb.12.117.

3. Kirkcaldy, R. D. et al. Neisseria gonorrhoeae Antimicrobial Susceptibility Surveillance — The Gonococcal Isolate Surveillance Project, 27 Sites, United States, 2014. MMWR. Surveillance Summaries 65, 1–19 (2016). DOI 10.15585/mmwr.ss6507a1.

4. World Health Organization. Global priority list of antibiotic-resistant bacteria to guide research, discovery, and development of new antibiotics. World Health Organization (2017).

5. Olesky, M., Zhao, S., Rosenberg, R. L. & Nicholas, R. A. Porin-Mediated Antibiotic Resistance in Neisseria gonorrhoeae: Ion, Solute, and Antibiotic Permeation through PIB Proteins with penB Mutations. Journal of Bacteriology 188, 2300–2308 (2006). DOI 10.1128/JB.188.7.2300-2308.2006.

6. Golparian, D., Shafer, W. M., Ohnishi,M. & Unemo, M. Importance of multidrug efflux pumps in the antimicrobial resistance property of clinical multidrug-resistant isolates of Neisseria gonorrhoeae. Antimicrobial agents and chemotherapy 58, 3556–9 (2014). DOI 10.1128/AAC.00038-14.

7. Sun, J., Deng, Z. & Yan, A. Bacterial multidrug efflux pumps: Mechanisms, physiology and pharmacological exploitations. Biochemical and Biophysical Research Communications 453, 254–267 (2014). DOI 10.1016/j.bbrc.2014.05.090.

8. Du, D., van Veen, H. W. & Luisi, B. F. Assembly and operation of bacterial tripartite multidrug efflux pumps. Trends in Microbiology 23, 311–319 (2015). DOI 10.1016/j.tim.2015.01.010.

9. Piddock, L. J. V. & Johnson, M. M. Accumulation of 10 fluoroquinolones by wild-type or efflux mutant Streptococcus pneumoniae. Antimicrobial agents and chemotherapy 46, 813–20 (2002). DOI 10.1128/AAC.46.3.813-820.2002.

10. Bohnert, J. A. et al. Site-directed mutagenesis reveals putative substrate binding residues in the Escherichia coli RND efflux pump AcrB. Journal of bacteriology 190, 8225–9 (2008). DOI 10.1128/JB.00912-08.

11. Fifer, H. et al. Failure of Dual Antimicrobial Therapy in Treatment of Gonorrhea. New England Journal of Medicine 374, 2504–2506 (2016). DOI 10.1056/NEJMc1512757.

12. Spratt, B. G. et al. Resistance of Neisseria gonorrhoeae to antimicrobial hydrophobic agents is modulated by the mtrRCDE efflux system. Microbiology 141, 611–622 (1995). DOI 10.1099/13500872-141-3-611.

13. Lei, H.-T. et al. Crystal structure of the open state of the Neisseria gonorrhoeae MtrE outer membrane channel. PloS one 9, e97475 (2014). DOI 10.1371/journal.pone.0097475.

14. Benz, R., Janko, K., Boos, W. & Läuger, P. Formation of large, ion-permeable membrane channels by the matrix protein (porin) of Escherichia coli. Biochimica et Biophysica Acta (BBA) - Biomembranes 511, 305–319 (1978). DOI 10.1016/0005-2736(78)90269-9.

15. Dargent, B., Hofmann,W., Pattus, F. & Rosenbusch, J. P. The selectivity filter of voltage-dependent channels formed by phosphoporin (PhoE protein) from E. coli. The EMBO journal 5, 773–8 (1986).

16. Andersen, C., Hughes, C. & Koronakis, V. Electrophysiological behavior of the TolC channel-tunnel in planar lipid bilayers. The Journal of membrane biology 185, 83–92 (2002). DOI 10.1007/s00232-001-0113-2.

17. Zachariae, U. et al. β-Barrel mobility underlies closure of the voltage-dependent anion channel. Structure 20, 1540–1549 (2012). DOI 10.1016/j.str.2012.06.015.

18. Nikaido, H. & Vaara, M. Molecular basis of bacterial outer membrane permeability. Microbiological reviews 49, 1–32(1985). DOI 10.1128/MMBR.67.4.593-656.2003.

19. Hanke, W. & Schule, W.-R. Physical properties of biological membranes and planar lipid bilayers. In Planar Lipid Bilayers (1993).

20. Surrey, T. & Jähnig, F. Refolding and oriented insertion of a membrane protein into a lipid bilayer. Proceedings of the National Academy of Sciences of the United States of America 89, 7457–7461 (1992). DOI 10.1073/pnas.89.16.7457.

21. Brunen, M. & Engelhardt, H. Asymmetry of orientation and voltage gating of the Acidovorax delafieldii porin Omp34 in lipid bilayers. European Journal of Biochemistry 212, 129–135 (1993). DOI 10.1111/j.1432- 1033.1993.tb17642.x.

22. Andersen, C., Koronakis, E., Hughes, C. & Koronakis, V. An aspartate ring at the TolC tunnel entrance determines ion selectivity and presents a target for blocking by large cations. Molecular Microbiology 44, 1131–1139 (2002). DOI 10.1046/j.1365-2958.2002.02898.x.

23. Wong, K. K., Brinkman, F. S., Benz, R. S. & Hancock, R. E. Evaluation of a structural model of Pseudomonas aeruginosa outer membrane protein OprM, an efflux component involved in intrinsic antibiotic resistance. Journal of bacteriology 183, 367–74 (2001). DOI 10.1128/JB.183.1.367-374.2001.

24. Janganan, T. K. et al. Opening of the outer membrane protein channel in tripartite efflux pumps is induced by interaction with the membrane fusion partner. The Journal of biological chemistry 286, 5484–93 (2011). DOI 10.1074/jbc.M110.187658.

25. Aksimentiev, A. & Schulten, K. Imaging α-Hemolysin with Molecular Dynamics: Ionic Conductance, Osmotic Permeability, and the Electrostatic Potential Map. Biophysical Journal 88, 3745–3761 (2005). DOI 10.1529/biophysj.104.058727.

26. Schulz, R. & Kleinekathöfer, U. Transitions between closed and open conformations of tolc: the effects of ions in simulations. Biophysical journal 96, 3116–3125 (2009).

27. Kutzner, C. et al. Insights into the function of ion channels by computational electrophysiology simulations. Biochimica et Biophysica Acta (BBA) - Biomembranes 1858, 1741–1752 (2016). DOI 10.1016/j.bbamem.2016.02.006.

28. Fitzpatrick, A. W. P. et al. Structure of the macab-tolc abc-type tripartite multidrug efflux pump. Nature Microbiology 2 (2017).

29. Oliver Beckstein, Philip C. Biggin & Sansom*, M. S. P. A Hydrophobic Gating Mechanism for Nanopores (2001). DOI 10.1021/JP012233Y.

30. Rasaiah, J. C., Garde, S. & Hummer, G. Water in Nonpolar Confinement: From Nanotubes to Proteins and Beyond. Annual Review of Physical Chemistry 59, 713–740 (2008). DOI 10.1146/an-nurev.physchem.59.032607.093815.

31. Andersen, C. et al. Transition to the open state of the TolC periplasmic tunnel entrance. Proceedings of the National Academy of Sciences of the United States of America 99, 11103–8 (2002). DOI 10.1073/pnas.162039399.

32. Janganan, T. K. et al. Opening of the outer membrane protein channel in tripartite efflux pumps is induced by interaction with the membrane fusion partner. The Journal of biological chemistry 286, 5484–93 (2011). DOI 10.1074/jbc.M110.187658.

33. Janganan,T. K., Bavro, V. N., Zhang, L., Borges-Walmsley, M. I. & Walmsley, A. R. Tripartite efflux pumps: energy is required for dissociation, but not assembly or opening of the outer membrane channel of the pump. Molecular microbiology 88, 590–602 (2013). DOI 10.1111/mmi.12211.

34. Song, J., Minetti, C. A., Blake, M. S. & Colombini, M. Meningococcal PorA/C1, a channel that combines high conductance and high selectivity. Biophysical journal 76, 804–13 (1999). DOI 10.1016/S0006-3495(99)77244-9.

35. Kutzner, C., Grubmüller, H., de Groot, B. L. & Zachariae, U. Computational electrophysiology: the molecular dynamics of ion channel permeation and selectivity in atomistic detail. Biophysical journal 101, 809–17 (2011). DOI 10.1016/j.bpj.2011.06.010.

36. Zachariae, U., Helms, V. & Engelhardt, H. Multistep mechanism of chloride translocation in a strongly anion-selective porin channel. Biophysical journal 85, 954–62 (2003). DOI 10.1016/S0006-3495(03)74534-2.

37. Pothula, K. R., Solano, C. J. & Kleinekathöfer, U. Simulations of outer membrane channels and their permeability. Biochimica et Biophysica Acta (BBA)-Biomembranes 1858, 1760–1771 (2016).

38. Askoura, M., Mottawea, W., Abujamel, T. & Taher, I. Efflux pump inhibitors (EPIs) as new antimicrobial agents against Pseudomonas aeruginosa. The Libyan journal of medicine 6 (2011). DOI 10.3402/ljm.v6i0.5870.

39. Higgins,M. K. et al. Structure of the ligand-blocked periplasmic entrance of the bacterial multidrug efflux protein TolC. Journal of molecular biology 342, 697–702 (2004). DOI 10.1016/j.jmb.2004.07.088.

40. Abraham, M. J. et al. GROMACS: High performance molecular simulations through multi-level parallelism from laptops to supercomputers. SoftwareX 1, 19–25 (2015). DOI 10.1016/j.softx.2015.06.001.

41. Yesylevskyy, S. O. ProtSqueeze: Simple and Effective Automated Tool for Setting up Membrane Protein Simulations. Journal of Chemical Information and Modeling 47, 1986–1994 (2007). DOI 10.1021/ci600553y.

42. Wolf, M. G., Hoefling,M., Aponte-Santamaría, C., Grubmüller, H. & Groenhof, G. g_membed: Efficient insertion of a membrane protein into an equilibrated lipid bilayer with minimal perturbation. Journal of Computational Chemistry 31, 2169–2174 (2010). DOI 10.1002/jcc.21507.

43. Lindorff-Larsen, K. et al. Improved side-chain torsion potentials for the Amber ff99SB protein force field. Proteins 78, 1950–8 (2010). DOI 10.1002/prot.22711.

44. Lindorff-Larsen, K. et al. Systematic validation of protein force fields against experimental data. PloS one 7, e32131 (2012). DOI 10.1371/journal.pone.0032131.

45. Berger, O., Edholm, O. & Jähnig, F. Molecular dynamics simulations of a fluid bilayer of dipalmitoylphosphatidylcholine at full hydration, constant pressure, and constant temperature. Biophysical journal 72, 2002–13 (1997). DOI 10.1016/S0006-3495(97)78845-3.

46. Cordomí, A., Caltabiano, G. & Pardo, L. Membrane Protein Simulations Using AMBER Force Field and Berger Lipid Parameters. Journal of chemical theory and computation 8, 948–58 (2012). DOI 10.1021/ct200491c.

47. Cho, C. H., Singh, S. & Robinson, G. W. Understanding all of water’s anomalies with a nonlocal potential. Journal of Chemical Physics 107, 7979–7988 (1997). DOI 10.1063/1.475060.

48. Joung, I. S., Cheatham, T. E. & III. Determination of alkali and halide monovalent ion parameters for use in explicitly solvated biomolecular simulations. The journal of physical chemistry. B 112, 9020–41 (2008). DOI 10.1021/jp8001614.

49. Miyamoto, S. & Kollman, P. A. Settle: An analytical version of the SHAKE and RATTLE algorithm for rigid water models. Journal of Computational Chemistry 13, 952–962 (1992). DOI 10.1002/jcc.540130805.

50. Hess, B., Bekker, H., Berendsen, H. J. C. & Fraaije, J. G. E. M. LINCS: A linear constraint solver for molecular simulations. Journal of Computational Chemistry 18, 1463–1472 (1997). DOI 10.1002/(SICI)1096-987X(199709)18:121463::AID-JCC4>3.0.CO;2-H.

51. Bussi, G., Donadio, D. & Parrinello,M. Canonical sampling through velocity rescaling. The Journal of Chemical Physics 126, 014101 (2007). DOI 10.1063/1.2408420.

52. Berendsen, H. J. C., Postma, J. P. M., van Gunsteren, W. F., DiNola, A. & Haak, J. R. Molecular dynamics with coupling to an external bath. The Journal of Chemical Physics 81, 3684–3690 (1984). DOI 10.1063/1.448118.

53. Feenstra, K. A., Hess, B. & Berendsen, H. J. C. Improving efficiency of large time-scale molecular dynamics simulations of hydrogen-rich systems. Journal of Computational Chemistry 20, 786–798 (1999). DOI 10.1002/(SICI)1096-987X(199906)20:8786::AID-JCC53.0.CO;2-B.

54. Huang, J. & MacKerell, A. D. CHARMM36 all-atom additive protein force field: Validation based on comparison to NMR data. Journal of Computational Chemistry 34, 2135–2145 (2013). DOI 10.1002/jcc.23354.

55. Jorgensen, W. L., Chandrasekhar, J., Madura,J. D., Impey, R. W. & Klein, M. L. Comparison of simple potential functions for simulating liquid water. The Journal of Chemical Physics 79, 926–935 (1983). DOI 10.1063/1.445869.

56. Evans, D. J. & Holian, B. L. The Nose-Hoover thermostat. The Journal of Chemical Physics 83, 4069–4074 (1985). DOI 10.1063/1.449071.

57. Parrinello, M. & Rahman, A. Polymorphic transitions in single crystals: A new molecular dynamics method. Journal of Applied Physics 52, 7182–7190 (1981). DOI 10.1063/1.328693. arXiv:1011.1669v3.

58. Jo, S., Kim, T., Iyer, V. G. & Im, W. CHARMM-GUI: A web-based graphical user interface for CHARMM. Journal of Computational Chemistry 29, 1859–1865 (2008).DOI 10.1002/jcc.20945.

59. Waterhouse, A. M., Procter, J. B., Martin, D. M. A., Clamp, M. & Barton, G. J. Jalview Version 2-a multiple sequence alignment editor and analysis workbench. Bioinformatics 25, 1189–1191 (2009). DOI 10.1093/bioinformatics/btp033.

60. Larkin, M. et al. Clustal W and Clustal X version 2.0. Bioinformatics 23,2947–2948 (2007). DOI 10.1093/bioinformatics/btm404.

61. Martí-Renom, M. A. et al. Comparative Protein Structure Modeling of Genes and Genomes. Annual Review of Biophysics and Biomolecular Structure 29, 291–325 (2000). DOI 10.1146/annurev.biophys.29.1.291.

62. Eswar, N. et al. Comparative Protein Structure Modeling Using MODELLER. In Current Protocols in Protein Science, 2.9.1–2.9.31 (John Wiley & Sons, Inc., Hoboken, NJ, USA, 2007).

63. Woodier, J., Rainbow, R. D., Stewart, A. J. & Pitt, S. J. Intracellular Zinc Modulates Cardiac Ryanodine Receptor-mediated Calcium Release. The Journal of biological chemistry 290, 17599–610 (2015). DOI 10.1074/jbc.M115.661280.

